# Inflammation-related pathology in the olfactory epithelium: its impact on the olfactory system in psychotic disorders

**DOI:** 10.1101/2022.09.23.509224

**Authors:** Kun Yang, Yuto Hasegawa, Janardhan P Bhattarai, Jun Hua, Milan Dower, Semra Etyemez, Neal Prasad, Lauren Duvall, Adrian Paez, Amy Smith, Yingqi Wang, Yun-Feng Zhang, Andrew P. Lane, Koko Ishizuka, Vidyulata Kamath, Minghong Ma, Atsushi Kamiya, Akira Sawa

## Abstract

Smell deficits and neurobiological changes in the olfactory bulb (OB) and olfactory epithelium (OE) have been observed in schizophrenia and related disorders. The OE is the most peripheral olfactory system located outside the cranium, and is connected with the brain via direct neuronal projections to the OB. Nevertheless, it is unknown whether and how a disturbance of the OE affects the OB in schizophrenia and related disorders. Addressing this gap would be the first step in studying the impact of OE pathology in the disease pathophysiology in the brain. In this cross-species study, we observed that chronic, local OE inflammation with a set of upregulated genes in an inducible olfactory inflammation (IOI) mouse model led to a volume reduction, layer structure changes, and alterations of neuron functionality in the OB. Furthermore, IOI model also displayed behavioral deficits relevant to negative symptoms (avolition) in parallel to smell deficits. In first episode psychosis (FEP) patients, we observed a significant alteration in immune/inflammation-related molecular signatures in olfactory neuronal cells (ONCs) enriched from biopsied OE and a significant reduction in the OB volume, compared with those of healthy controls (HC). The increased expression of immune/inflammation-related molecules in ONCs was significantly correlated to the OB volume reduction in FEP patients, but no correlation was found in HCs. Moreover, the increased expression of human orthologues of the IOI genes in ONCs was significantly correlated with the OB volume reduction in FEP, but not in HCs. Together, our study implies a potential mechanism of the OE-OB pathology in patients with psychotic disorders (schizophrenia and related disorders). We hope that this mechanism may have a cross-disease implication, including COVID-19-elicited mental conditions that include smell deficits.

## Introduction

Smell deficits, measured by established psychophysical measures (odor identification, odor discrimination, and odor detection threshold), have been reproducibly reported in patients with psychotic disorders, such as schizophrenia and related disorders^1–8^. Smell deficits are also observed in patients with first episode psychosis (FEP)^3,9–11^. Importantly, the correlation of smell deficits in these severe mental disorders is tightly observed with negative symptoms and some domains of cognitive deficits, whereas rarely with positive symptoms^2,3,12–19^. Smell deficits also appear to predict poor outcomes in patients with schizophrenia and can identify patients at high risk of developing unremitting negative symptoms, particularly anhedonia^15,20^. Altogether, smell deficits are not merely a secondary outcome of confounding factors, such as medications.

Instead, smell deficits may be a reflection of specific pathophysiological mechanisms underlying schizophrenia and related disorders. Anatomically, olfactory sensory neurons (OSNs) in the olfactory epithelium (OE) directly project to the glomeruli in the olfactory bulb (OB), where they form synapses on mitral and tufted cells^21^. Mitral and tufted cells in the OB then project to the primary olfactory cortex, where the information from OSNs is transmitted to the higher cortex, such as the orbitofrontal cortex and medial prefrontal cortex, as well as other central brain regions that are involved in motivation, emotion, and cognition^22^. Studies in basic neurobiology have deciphered a mechanistic link between the OE and the brain circuitry in which the OB plays a crucial role as the gateway of higher brain circuitries at the functional levels^23,24^. Our group previously reported an inducible olfactory inflammation (IOI) mouse model in which induction of local and chronic OE inflammation switches the fate of neuroepithelial stem cells from neurogenesis to immune defense, with a set of inflammation-related genes (called IOI genes) upregulated.

The OE is the most peripheral olfactory system located outside the cranium. As a result, the OE is directly exposed to air pollution and viral infections in the upper respiratory tract. Intranasal infection with SARS-CoV-2 associated with smell deficits has frequently resulted in brain dysfunction, including the development of schizophrenia-related clinical manifestations^25,26^. Likewise, air pollution has been reproducibly reported as a major risk factor for schizophrenia and related disorders^27–29^. For subjects intrinsically vulnerable to mental illnesses, the OE can be a nidus of interactions between genetic and environmental risk factors for the disease. Molecular and cellular changes in the OE in patients with schizophrenia and related disorders have been reported from multiple groups^30–38^.

One outstanding question is whether and how the OE pathology may impact the brain pathophysiology in patients with schizophrenia and related disorders. Given that the OSNs in the OE directly project to the OB, which functions as the gateway for olfactory-higher brain neurocircuits (e.g., the olfactory-prefrontal circuits), defining the OE pathology and its impact on the OB will be the crucial first step to address this question. To address this knowledge gap, we hypothesized that inflammation-related pathological processes exist in the OE of FEP patients, given that the OE is a tissue where psychosis-associated genetic and environmental factors come across. We assumed that olfactory neuronal cells (ONCs) enriched from the nasal cavity via biopsy^39^, represented pathological signatures at the molecular level. We further hypothesized that a mouse model with OE local inflammation, such as the IOI mouse model, might mimic the pathological changes in the OE found in FEP patients.

In the present study, we showed that chronic local OE inflammation led to anatomical and layer structure changes in the OB, reduced synaptic inputs to OB neurons, and behavioral deficits relevant to negative symptoms (avolition) in parallel to smell deficits, in the IOI model. We also observed immune/inflammation-related molecular signatures in ONCs and a significant volume reduction of the OB in FEP patients compared with healthy controls (HCs). Importantly, the increased expression of immune/inflammation-related molecules in ONCs was significantly correlated to the reduction of the OB volume in FEP patients, but such a correlation was not observed in HCs. Moreover, the increased expression of the human orthologues of the IOI genes in ONCs was also significantly correlated with the reduction of the OB volume in FEP patients. No such a correlation was observed in HCs. Together, these results suggest an association between OE inflammation and OB pathology in schizophrenia and related disorders.

## Materials and Methods

### IOI mouse model

The IOI mouse model was generated by crossing mice expressing an exogenous TNF-α transgene under the Tet-response element (TRE) with a strain carrying the reverse tetracycline transcriptional activator (rtTA) under the control of the Cyp2g1 promoter, which has been standardized in the C57BL/6 background as previously described^40–42^. For anatomical assessment of the OB, three-to four-month-old IOI mice were treated with 0.2g/kg doxycycline (DOX) containing food for 6 weeks to induce TNF-α expression in the sustentacular cells of the OE, leading to chronic and local OE inflammation. To examine the effect of OE inflammation on behaviors in young adulthood, one month-old IOI mice (DOX-treated *Cyp2g1-rtTA;TRE-TNF*) and controls (DOX-treated *TRE-TNF* single-transgenic mice) of both sexes were fed a DOX-containing diet for 30-40 days, followed by behavioral experiments.

Both male and female mice were used in each experiment. The sample size for each experiment was determined using the pilot result. A power level of 0.80 and significance level of 0.05 were selected. The animals in the experiments were littermates, and used the same age and sex. Mice were first categorized based on their genotype, and then randomly distributed among the experimental groups. All data collection and analysis were conducted by experimenters who were blinded to the groups. All animal procedures were approved by the Institutional Animal Care and Use Committee of Johns Hopkins University School of Medicine, and adhered to ethical considerations in animal research.

### Anatomical assessment of the OB in the IOI model

Mouse brains were extracted after perfusion with 4% paraformaldehyde (PFA). The fixed brains were embedded in cryocompound (Sakura Finetek) after replacement of PFA with 30% sucrose in phosphate buffered saline. Coronal sections were obtained at 20 µm with a cryostat (Leica). The length and width of the OB was measured as shown in **Figure 1A**.

**Figure 1.**
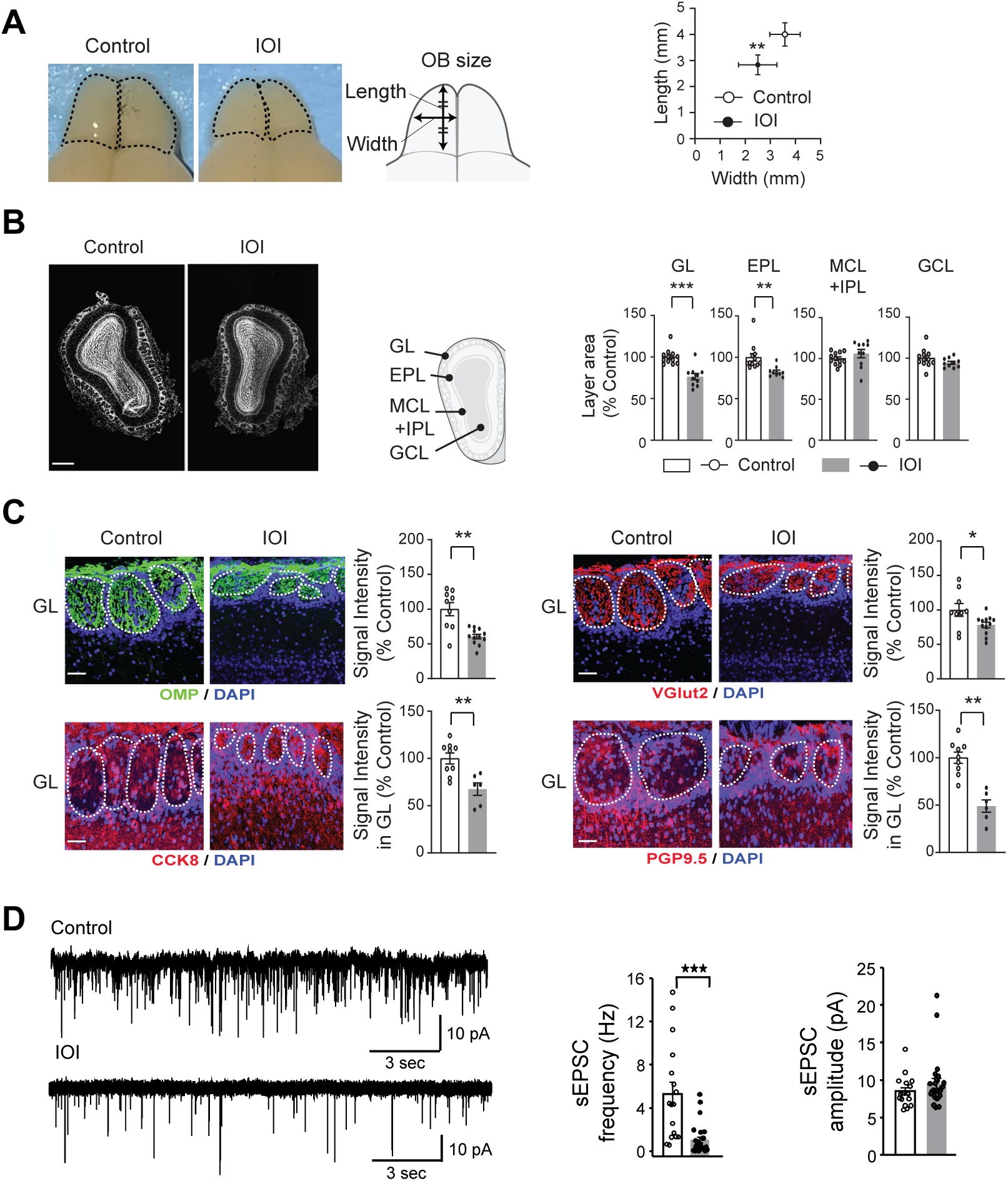
Structural alterations in the OB of the IOI mice. Abbreviations: GL, glomerular layer; EPL, external plexiform layer; MCL, mitral cell layer; IPL, internal plexiform layer; GCL, granule cell layer. **(A)** OB size (*n* = 6 mice per condition): left panel: dorsal view of the OB, and schematic diagram of the OB measurement methods; right panel: quantification of the OB length and width. **(B)** DAPI staining of the OB (white) (*n* = 10 mice per condition): left panel: representative coronal section image of the OB. Scale bar, 500 μm; middle panel: schematic diagram of the OB coronal section layers; right panel: quantification of each area as % of control mice. **(C)** Immunohistochemistry with antibody against OMP (green), VGlut2 (red), CCK8 (red), and PGP9.5 (red) with DAPI staining in the GL (*n* = 6-12 mice per condition): left panel: representative images of the GL. Scale bar, 50 μm; right panel: quantification of the signal intensity of OMP, VGlut2, CCK8, and PGP9.5 as % of control. **(D)** Patch clamp recording of spontaneous EPSCs from OB tufted cells: left panel: raw traces; middle panel: averaged frequency; right panel: amplitude. For all cells, the holding potential was -70 mV under voltage clamp mode. \*\*\**p* < 0.001, \*\**p* < 0.01, \**p* < 0.05.

The OB layer structure was further examined using DAPI signal to divide the middle section of the OB into layers. The area of each layer was then measured using ImageJ-FIJI software (https://imagej.net/Fiji). Signal intensity was also measured by creating a 2D mask of the affected area and examining the mean fluorescence of signal in all sections using ImageJ-FIJI software. The same imaging parameters and thresholds were used for both IOI and control mice to minimize the experimental bias.

### Immunohistochemistry in the IOI model

Immunohistochemistry was performed using our previously published methods with some modifications^43^. Briefly, brain sections were heated in HistoVT One solution for 30 min at 60°C for antigen retrieval, followed by blocking procedures. Sections were then incubated with primary antibodies at 4°C overnight. The primary antibodies include an anti-OMP antibody (Wako, 1:1000), an anti-VGlut2 antibody (Sigma-Aldrich, 1:100), an anti-PGP9.5 antibody (NOVUS, 1:200), and an anti-CCK8 antibody (Sigma-Aldrich, 1:500). After this, sections were incubated for 2 h with secondary antibodies conjugated with Alexa 488 (Invitrogen, 1:400), Alexa 568 (Invitrogen, 1:400), and Alexa 647 (Invitrogen, 1:400). Lastly, nuclei were labeled with DAPI (Roche). Immunofluorescence images were acquired using the Zeiss LSM700 confocal microscope with ZEN 2010 software.

### Behavioral assessment in the IOI model

After DOX treatment followed by 1 week habituation for reversed 12-h light/dark cycle, during the dark period of the cycle, we conducted the following tests.

#### Open field test

Following our established protocol^43^, the locomotor activity was assessed over a 30-min timeframe in 40 x 40 cm activity chambers equipped with infrared beams (PAS system, San Diego Instruments). Any horizontal and vertical locomotor activity that occurred in the center or along the periphery walls was automatically captured by the beam breaks.

#### Olfactory habituation/dis-habituation test

Olfaction was evaluated using the olfactory habituation/dis-habituation test, following our established protocol^40^. The odors were presented to the mice via a suspended cotton swab placed inside a clean cage filled with fresh shavings. Each mouse underwent three consecutive 2-min trials for each odor, with 2-min intervals between trials. The duration during which the mouse sniffed the swab was recorded. Three odors were tested in this study: unscented water, vanilla (1:33; GEL SPICE), and banana (1:100; McCormick).

#### Progressive ratio (PR) schedule of reinforcement task

The PR schedule task assessed the progressively increasing number of responses needed before a reward was given during a PR test session. We quantified the motivation for the food reward using the “breakpoint”, the amount of lever pressing needed for the reward. The animals were individually housed and habituated to a reversed light/dark cycle. Food intake was restricted until the target weight (80-90% of the *ad libitum* feeding weight) was achieved. Then the PR training started as follows: mice were placed in an operant chamber box (7.5 in x 7.5 in x 10 in) equipped with a food delivery port, 2 levers, and a tone generator (PACKWIN system, Harvard Apparatus). The task was administered under red light. First, fixed-ratio tasks were conducted. Mice could initially get food reward, a 20 mg sugar pellet, by pressing the lever once. Once the target (1 session of 50 rewards or 3 sessions of more than 30 rewards) was achieved during a 1 h session, tasks were transitioned to 3 presses, and then 5 presses for reward delivery. Then, progressive-ratio tasks were administered. Mice had to press the lever an increasing number of times (0, 1, 2, 4, 6, 9, 12, 15, 20, 25, 32, 40, 50, 62, 77, 95, 118, 145, 178, 219, 268) over 2 h to earn a food pellet. The number of total lever presses, total earned food pellets, and final breakpoint reached were recorded in the 2 h test session.

#### Free feeding test

According to published protocol^44^, each mouse was subjected to an 18-h period of fasting, then was provided with a single food pellet of standard lab chow or sugar pellet in their home cage. Their free-feeding behavior was observed for 60 min. The weight of the consumed food (in grams) was then measured and recorded.

#### Elevated plus maze test

Following our published protocol^43^, a mouse was positioned at the central intersection of the arms within the plus maze (San Diego Instruments) and observed through videography for 5 min. The frequency of entries into both the closed and open arms, as well as the time spent in each arm, was recorded. The average percentage of entries into the open arms was calculated.

### Patch clamp recording in mouse brain slices

Mice were deeply anesthetized with ketamine-xylazine (200 and 15 mg/kg body weight, respectively) and decapitated. The brain was dissected out and immediately placed in ice-cold HEPES buffer. Coronal OB slices (130 μm) were cut using a Leica VT 1200S vibratome. Slices were incubated in oxygenated artificial cerebrospinal fluid (ACSF) containing (in mM): 126 NaCl, 2.5 KCl, 2.4 CaCl2, 1.2 MgSO4, 11 D-glucose, 1.4 NaH2PO4 and 25 NaHCO3 (osmolality ∼305 mOsm and pH 7.4, bubbled with 95% O2−5% CO2) for 1 h at 31°C and kept in oxygenated ACSF at room temperature thereafter. Before recording, slices were transferred to a recording chamber and continuously perfused with oxygenated ACSF. Recording pipettes were made from borosilicate glass (GC210F-10; Harvard Apparatus) with a Flaming-Brown P-97 puller (Sutter Instruments; tip resistance 5–8 MΩ). The pipette solution contained (in mM): 120 K-gluconate, 10 NaCl, 1 CaCl2, 10 EGTA, 10 HEPES, 5 Mg-ATP, 0.5 Na-GTP, and 10 phosphocreatine. Electrophysiological recordings were controlled by an EPC-10 amplifier combined with Pulse Software (HEKA Electronic) and analyzed using Igor Pro and mini-analysis (Synaptosoft, Inc.).

### Human study participants

The present study with human subjects was conducted in accordance with The Code of Ethics of the World Medical Association (1964 Declaration of Helsinki) and was approved by the Johns Hopkins School of Medicine Institutional Review Board. Written informed consent was obtained for all participants.

Adolescents and young adults between 18 and 35 years old were recruited from within and outside of the Johns Hopkins Hospital. All the subjects were recruited before the COVID pandemic. Exclusion criteria included history of traumatic brain injury, neurologic condition, intellectual disability, cancer, viral infection, active substance abuse, and conditions affecting olfaction (e.g, history of sinus surgery, rhinoplasty, chronic rhinosinusitis). Controls were screened and excluded for a family history of psychotic disorders. Patients must be within 24 months of the onset of psychotic manifestations as assessed by study team psychiatrists using the Structured Clinical Interview for DSM-IV (SCID) and information from available medical records.

The present study investigated the data from 89 HCs and 93 FEP patients. Data from all these subjects were used for the OB volume assessment. These patients included those with schizophrenia (*n* = 50), schizoaffective disorder (*n* = 10), schizophreniform disorder (n=3), bipolar disorder with psychotic features (*n* = 21), major depressive disorder with psychotic features (*n* = 5), brief psychotic disorder (n=2), and psychotic disorder not otherwise specified (n=2). A demographic summary of these 182 (89 HCs and 93 FEP patients) subjects is shown in **Table 1A**. All the demographic variables were adjusted in data analyses (see details in “Statistical analysis”).

**Table 1.** Demographics of the study cohorts. Significant results are highlighted in bold. Abbreviations: SD, standard deviation; Y, yes; N, no; AA, African American.

**(A) Entire Cohort: subjects with OB assessments**
|  | **HCs (n=89)** | **FEP patients (n=93)** | **p-value** |
| --- | --- | --- | --- |
| Age (mean/SD) | 23.53 / 3.89 | 22.11 / 4.44 | **0.02** |
| Sex (Male/Female) | 40 / 49 | 68 / 25 | **2.02E-04** |
| Race (AA/Caucasian/Asian/other) | 56 / 25 / 2 / 6 | 45 / 34 / 5 / 9 | 0.07 |
| Tobacco use (Y/N) | 3 / 86 | 24 / 69 | **5.17E-05** |

**(B) Subset Cohort: subjects with both OB assessments and RNA-Seq from ONCs**
|  | **HCs (n=59)** | **FEP patients (n=44)** | **p-value** |
| --- | --- | --- | --- |
| Age (mean/SD) | 23.33 / 3.25 | 22.75 / 3.88 | 0.42 |
| Sex (Male/Female) | 31 / 28 | 30 / 14 | 0.16 |
| Race (AA/Caucasian/Asian/other) | 37 / 16 / 2 / 4 | 26 / 14 / 1 / 3 | 0.87 |
| Tobacco use (Y/N) | 2 / 57 | 8 / 36 | **0.03** |

**(C) Comparison between the entire and subset cohorts**
|  | **p-value**<br>**(89 HCs vs. 59 HCs)** | **p-value**<br>**(93 FEPs vs. 44 FEPs)** |
| --- | --- | --- |
| Age | 0.75 | 0.40 |
| Sex | 0.46 | 0.69 |
| Race | 1.00 | 0.32 |
| Tobacco use | 1.00 | 0.44 |
| Total OB | 0.75 | 0.99 |
| Right OB | 0.87 | 0.99 |
| Left OB | 0.64 | 1.00 |

Among 89 HCs and 93 FEP patients whom we assessed for the OB volume, we collected ONCs from 59 HCs and 44 FEP patients (20 schizophrenia, 6 schizoaffective disorder, 2 schizophreniform disorder, 12 bipolar disorder with psychotic features, and 3 major depressive disorder with psychotic features, and 1 psychotic disorder not otherwise specified). A demographic summary of these 103 subjects (59 HCs + 44 FEP patients) is shown in **Table 1B**. Notably, we didn’t find any significant difference in the demographics of HCs between the entire (89 HCs) and subset (59 HCs with ONCs) cohorts. Likewise, we did not find any significant difference in the demographics of FEPs between the entire (93 FEP patients) and subset (44 FEP patients with ONCs) cohorts (**Table 1C**). These results suggested that there was no cohort bias between these two cohorts.

### ONCs obtained via nasal biopsy from human subjects

Nasal biopsy followed by the enrichment and collection of ONCs was conducted according to our established protocol shown in past publications^35,38,39,45^. Briefly, nasal biopsied tissues were collected under endoscopic control with local anesthesia and dissociated by mechanical and enzymatic treatments. After removing tissue debris, cells were plated on 6-well plates (Day 0 plate). Cells floating or loosely attached to the surface were transferred from the Day 0 plate to a Day 2 plate and from a Day 2 plate to a Day 7 plate. Through these processes, connective tissue-origin cells and non-neuronal cells that are attached to Day 0 and Day 2 plates are removed. Accordingly, in the present study, we examined cells in Day 7 plates called ONCs. ONCs were stored in liquid nitrogen tanks for further experiments.

### Molecular expression profiles (RNA-Seq) of ONCs

Total RNA was isolated from ONCs using the RNeasy Plus Mini Kit (Qiagen). RNA quality was assessed on the Agilent Fragment Analyzer using an RNA High Sensitivity kit (DNF-472) and quantified using a Qubit 4 RNA BR kit (Thermo Fisher). RNA libraries were prepared with 500 ng total RNA. Library generation was accomplished using the NEBNext Ultra II Directional RNA Library Prep Kit for Illumina (E7760 and E7490) following the NEBNext Poly(A) mRNA Magnetic Isolation Module protocol. Libraries were enriched using 11 cycles of PCR amplification. Library quality and quantification was assessed on the Agilent Fragment Analyzer using a High Sensitivity NGS Kit (DNF-474) and a Qubit 4 RNA BR kit (Thermo Fisher). Samples were then normalized to 4 nM and pooled in equimolar amounts. Paired-End Sequencing was performed using Illumina’s NovaSeq6000 S4 200 cycle kit.

### OB volume measurement in human subjects

All images were acquired on a 3T Philips MRI scanner (Philips Healthcare, Best, The Netherlands). The three-dimensional T2-weighted scan was acquired for each participant with the following parameters: voxel = 1.1×1.1×2.2 mm^3^, 70 slices, repetition time (TR) = 4165 ms, echo time (TE) = 80 ms, flip angle = 90°, acquisition time = 3 minutes 28 seconds.

The FMRIB Software Library (FSL, https://fsl.fmrib.ox.ac.uk/fsl/) software was used to normalize the T2-weighted images into the MNI space. Subsequent analysis was performed in Insight ToolKitSNAP (ITK-SNAP) (www.itksnap.org). The OB was manually identified and segmented on coronal images, with additional corrections made in the sagittal and axial planes. Volumetric measures were obtained using the built-in functions in ITK-SNAP at the level of the anterior cribriform plate following the procedures established previously^46^. Two experienced researchers (NP and LD) performed the segmentation separately and were blinded to participant information. After all segmentation was completed, discrepancies between the two researchers were assessed and final measurements were agreed upon. The intraclass correlation coefficient for inter-rater reliability between raters (NP and LD) was 0.86.

### Statistical analysis

R version 4.1.2 was used to conduct statistical analysis. For the preclinical study, t-test was conducted to compare data between the IOI mice and control mice. For the clinical study, t-test and chi-squared test were conducted to compare demographics between groups for continuous and categorical variables, respectively. Most of the patients, except five patients, were medicated. Antipsychotic medication dosages were converted to chlorpromazine (CPZ) equivalents using published reference tables with the Defined Daily Doses (DDDs) method^47^.

#### RNA-seq analysis in human ONCs

Data preprocessing: FastQC^48^ was used to check the quality of reads. High-quality data were obtained from raw data by using cutadapt^49^ to remove adapters, primers, and reads with low quality (option -q 10) or shorter than 20 nt. Hisat2 (option --dta)^50^ was used to map the clean reads to the human genome, version GRCh38 (Genome Reference Consortium Human Build 38). Stringtie^50^ was used to assemble and merge transcripts and estimate transcript abundance. A Python script (prepDE.py) provided by the Stringtie developer^50^ was used to create count tables for further analysis. Principle component analysis (PCA) was conducted for quality control and no outliers were detected in our dataset.

##### Differential expression analysis

R library *DESeq2*^51^ was utilized to compare the expressional profiles between FEP patients and HCs. Age, sex, race, tobacco usage status (Yes/No), and one hidden/unknown confounding factor identified by *sva*^52^ were included as covariates in the design formula. The Benjamini and Hochberg procedure was used for multiple comparison correction. Genes with false discovery rates (FDR) smaller than 0.05 were considered significant.

##### Co-expression gene network analysis

Linear regression was conducted to calculate the adjusted transcripts per million (TPM) values controlling for age, sex, race, tobacco usage status, and one hidden/unknown confounding factor identified by *sva*^52^. A weighted correlation network analysis (WGCNA)^53^ was then conducted to construct co-expression gene networks using adjusted TPM values, and the expression of the identified network module(s) was compared between FEP patients and HCs via two-tailed analysis of covariance (ANCOVA) controlling for age, sex, race, tobacco usage status and one hidden/unknown confounding factor identified by *sva*^52^. The Benjamini and Hochberg procedure was used for multiple comparison correction. Lastly, the biological functions of network modules were examined via functional enrichment analysis using *gProfiler*^54^. Additionally, the potential involvement of genes in immune/inflammation-related processes was assessed by literature mining of PubMed. We searched PubMed using each gene with the keywords “immune,” “inflammation,” or “inflammatory“. A gene that appeared with either keyword in more than 20 publications was defined as immune/inflammation-related. We also tested the robustness of our results via additional analyses using more than 30 or 40 publications as the threshold to define the immune/inflammation-related genes.

#### OB volume assessment in human subjects

Two-tailed ANCOVA with age, sex, race, tobacco usage status, and intracranial volume as covariates was performed to compare the total, left, and right OB volumes between FEP patients and HCs.

#### Association between OE inflammation and OB volume

When we looked for an association between immune/inflammation-related differentially expressed genes (DEGs) and the OB volume, PCA analysis was conducted for the matrix of expression levels of immune/inflammation-related DEGs. The first principal component was used to measure the total expression of immune/inflammation-related DEGs. Two-tailed partial correlation analysis controlling for age, sex, race, tobacco usage status, intracranial volume, duration of illness, and CPZ dose was then conducted to examine whether an increase in immune/inflammation-related DEGs was correlated with a reduction of the OB volume in FEP patients.

When we integrated the mouse and human data to further explore the association between OE inflammation and OB volume, we conducted PCA analysis and used the first principal component to measure the total expression of the human orthologues of the IOI genes^41^ in ONCs. We then examined the correlation between the total expression and the OB volume in FEP patients, via two-tailed partial correlation analysis controlling for age, sex, race, tobacco usage status, intracranial volume, duration of illness, and CPZ dose.

## Results

### Mouse study: pathological impact of chronic, local OE inflammation on the OB anatomy, OB volume, and behaviors in the IOI model

We examined the impact of chronic, local OE inflammation on the OB volume and structure in young adult male and female mice. After exposure to local OE inflammation for 6 weeks (starting when mice were 3-4 months old in young adulthood), we observed significant reductions in the length (*p* = 9.87E-03) and width (*p* = 3.65E-03) of the OB in the IOI mice compared with control mice (**Figure 1A**). We next examined the influence of OE inflammation on the layer structure of the OB. The glomerular layer (*p* = 1.18E-07) and external plexiform layer (EPL) (*p* = 3.30E-05) in the OB of the IOI mice were smaller than those of the control mice. In contrast, no significant differences between the two groups were observed in the granule cell layer and the layer that included both the mitral cell and internal plexiform layers (**Figure 1B**).

At the immunohistochemical level, we observed a reduction in the expression of OMP (a marker for mature OSNs), CCK8 (a marker for tufted cells), and PGP9.5 (a marker for mitral cells) in the glomerular layer of the IOI model (**Figure 1C**). Furthermore, we also observed a reduction in Vglut2 (a glutamatergic pre-synapse marker) staining in the glomerular layer of the IOI model (**Figure 1C**). These data suggest that local OE inflammation elicits a reduction in OSNs, which would lead to reduced synaptic connectivity between OSNs and OB neurons.

To directly test this hypothesis, we performed patch clamp recordings on OB neurons in acute brain slices. We targeted tufted cells in the external plexiform layer because they receive strong excitatory inputs from OSNs and can be readily identified based on size, morphology and firing patterns^55^. With a holding potential at -70 mV under the voltage clamp mode, which minimizes the contribution of GABA_A_-mediated inhibitory currents, we found a significant reduction in the frequency of spontaneous excitatory postsynaptic currents (sEPSCs) in the neurons from the IOI mice compared to controls (**Figure 1D**). The glomerular size in the IOI mice is much smaller than that of the control mice (**Figure 1B**), and the glomerular size is linearly correlated with the number of input OSNs^56^. Taken together, although we can’t completely rule out that an enhanced network inhibition might influence the release probability from OSNs, the most likely mechanism indicated by these data is a reduction of OSN synaptic inputs to tufted cells. In contrast, there was no significant change in the amplitude of sEPSCs, suggesting normal glutamate receptor signaling in postsynaptic cells in the IOI mice. Taken together, our data show that chronic, local OE inflammation results in structural and functional changes in the OB.

We previously reported smell deficits in the IOI model by through electro-olfactogram recordings and olfactory habituation/dis-habituation test^40,42^. In the present study, through olfactory habituation/dis-habituation test, we confirmed the presence of smell deficits in the IOI mice (**Figure 2A**). A strong association between smell deficits and negative symptoms has been reproducibly reported in patients with psychotic disorders^2,3,12–19^. Thus, we performed PR schedule of reinforcement task, a representative behavioral assay relevant to negative symptoms (particularly associated with avolition) using the same cohort of the IOI mice and controls (**Figure 2B**). Both IOI and control groups had similar increases in the lever pressing over time in the fixed ratio reinforcement training sessions. In the PR test session, the IOI group displayed a lower number of total lever presses (*p* = 4.99E-05) and a reduced number of earned food pellets (*p* = 7.19E-04), compared to controls (**Figure 2C**). These data were consistent with the reduction in the breakpoint of the IOI mice (a decrease of 44.9%; *p* = 5.36E-04) (**Figure 2C**). To exclude the possibility that the PR test results were affected by smell deficits, we further conducted the free feeding behavioral test. the IOI mice exhibited normal food intake under free-access conditions and displayed a preference for high-sugar food pellets over regular food pellets at the physiological level (**Figure 2D**), indicating that smell deficits are unlikely to influence the outcome of the PR task. In addition, no difference was observed in body weight between the IOI and control mice (**Figure S1A**). No deficits in open field behaviors and elevated plus maze tests (**Figure S1B, S1C**). Taken together, chronic, local OE inflammation in young adulthood leads to avolition-like behavioral deficits in parallel to smell deficits.

**Figure 2.**
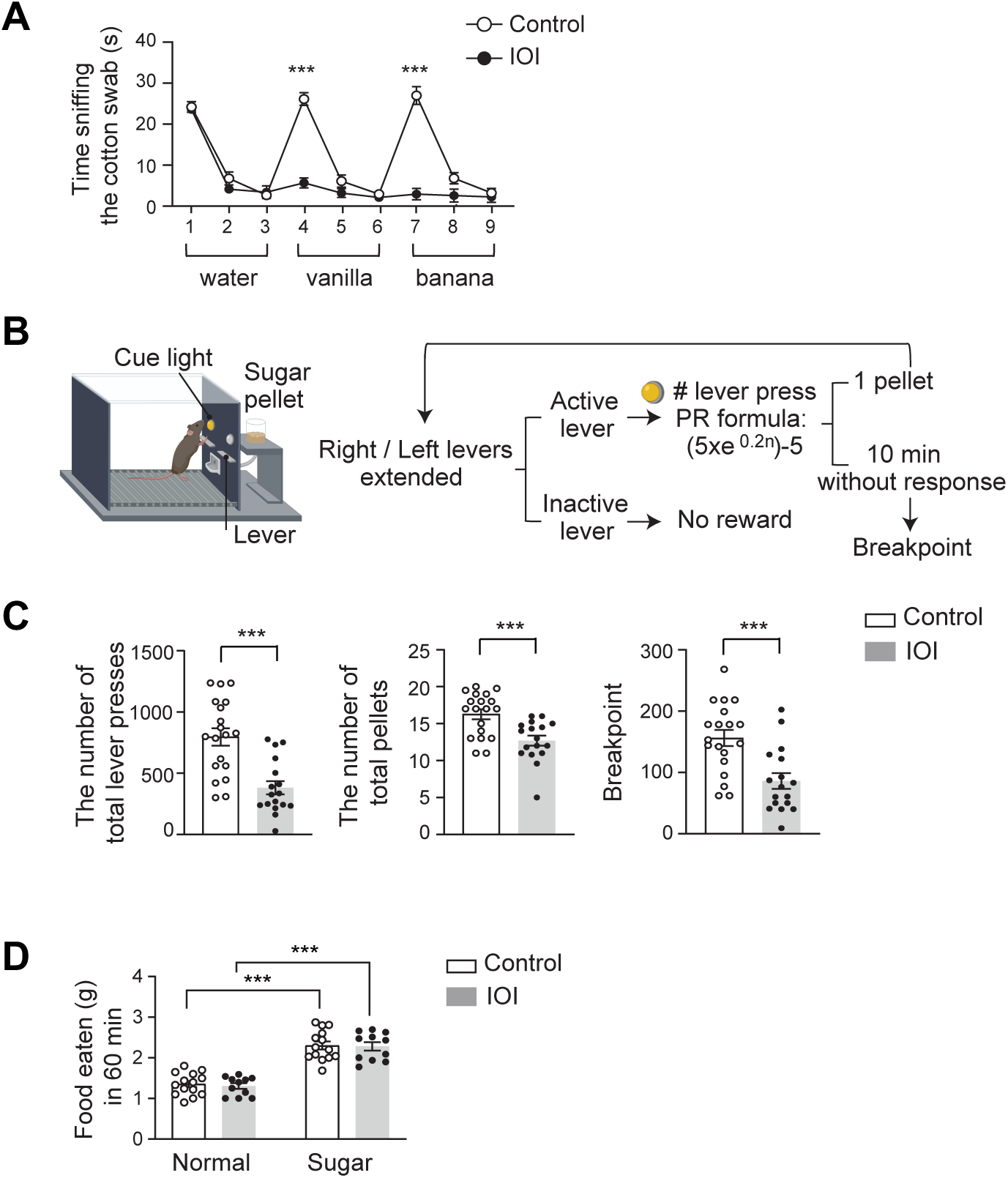
Behavioral deficits in the IOI mice. **(A)** Olfactory habituation/dis-habituation test. The mice were subjected to three consecutive exposures to water and two distinct odorants, such as vanilla and banana. Control, n=9 (4 males, 5 females); IOI, n=7 (4 males, 3 females) **(B)** The schema of PR schedule of reinforcement task using an operant conditioning chamber. **(C)** Total number of lever presses, pellets received, and animal’s breakpoints in PR test session were shown. Control, n=19 (9 males, 10 females); IOI, n=17 (8 males, 9 females). **(D)** Free feeding test. The weight of the consumed standard lab chow (normal food pellet) or high-sugar pellet food in 60 min was measured under free-feeding conditions. Control, n=14 (6 males, 8 females); IOI, n=11 (8 males, 3 females). \*\*\**p* < 0.001.

### Human study: immune/inflammatory molecular signature in OE-derived neuronal cells (ONCs) from patients

Previously, we reported preliminary observations of altered molecular signatures in ONCs from FEP patients (n=16) compared with HCs (n=22)^57^. In the present study, we expanded the sample size (44 FEP patients and 59 HCs: see **Table 1B**) and conducted a more comprehensive analysis, particularly testing our hypothesis that immune/inflammatory molecular changes exist in the OE-derived neuronal cells from FEP patients.

We conducted a genome-wide differential expression analysis to identify genes that were differentially expressed in FEP patients compared with HCs, while controlling for confounding factors and correcting for multiple comparisons. Our analysis identified 27 DEGs (FDR < 0.05) in FEP patients (**Table S1**). We then conducted literature mining of PubMed by searching for publications using each gene with the keyword “immune,” “inflammation,” or “inflammatory.” Here a gene that appeared with any keyword in more than 20 publications was defined as immune/inflammation-related. Based on these criteria, 33.3% of the DEGs (9 out of 27) were categorized as immune/inflammation-related, whereas 24.5% of the non-DEGs (3,413 out of 13,908) were categorized as immune/inflammation-related. Notably, this higher proportion of immune/inflammation-related genes in DEGs compared to non-DEGs was consistent even when we changed the threshold (the number of publications) to define the immune/inflammation-related genes: 30 publications as the threshold: 29.6% of DEGs (8 out of 27) and 19.7% of non-DEGs (2,738 out of 13,908); 40 publications as the threshold: 25.9% (7 out of 27) of DEGs and 16.7% of non-DEGs (2,323 out of 13,908). These findings suggest that FEP patients have altered immune/inflammation-associated molecular signatures in ONCs.

We further conducted a co-expression gene network analysis. Six co-expression network modules were identified, and 2 out of these 6 network modules were significantly altered in FEP patients compared with HCs: module I includes 124 genes, and module II includes 223 genes (**Table S2**). We then conducted a functional enrichment analysis to examine the biological functions of each network module. We found that genes related to the nervous system development and the immune system process were enriched in module II, while genes related to the metabolic process of serine and glutamate were enriched in module I. A further PubMed search found that 31.8% of genes in module II (71 out of 223) appeared in more than 20 publications with the keywords “immune,” “inflammation,” or “inflammatory” (**Table S3**). In contrast, only 24.4% of non-module II genes (3,351 out of 13,712) were immune/inflammation-related. In summary, we observed a higher proportion of immune/inflammation-related genes in module II than in other modules. These data also endorse the notion that FEP patients have immune/inflammation-related molecular signatures in ONCs, which likely reflect OE inflammation.

### Human study: reduced OB volume in FEP patients

Our group has reported preliminary observations of a reduced OB volume derived from T1-weighted (T1w) images at 3 Tesla in FEP patients (n=16) compared with HCs (n=22)^57^. In the present study, we addressed the OB volume more systematically with T2w images in a much larger sample size: 93 FEP patients and 89 HCs (**Table 1A**). Compared with T1w images, T2w images are more precise in assessing the OB volume due to a brighter signal of the cerebrospinal fluid, which helps define the edges of the OB more clearly (**Figure 3A**).

**Figure 3.**
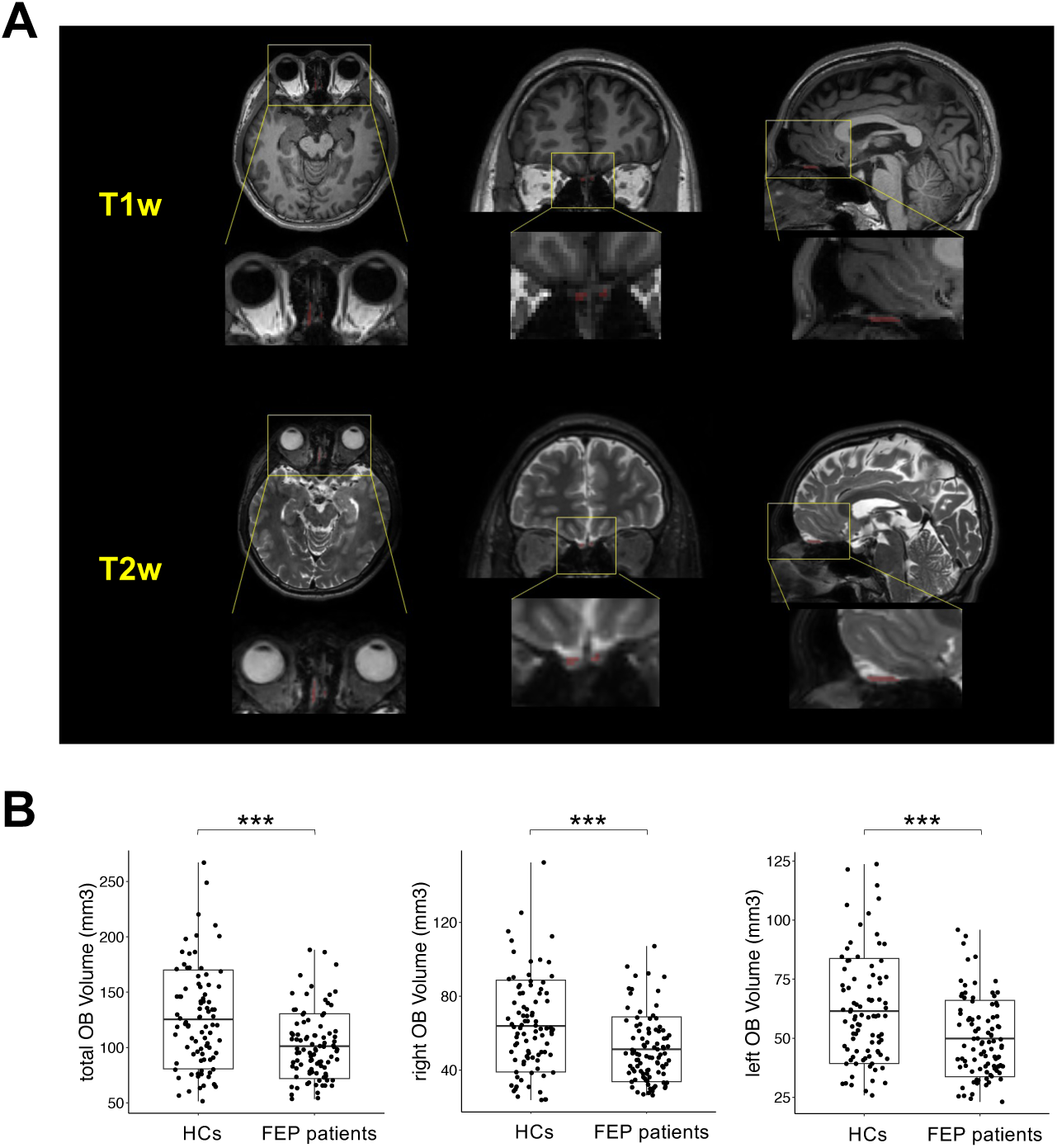
Volumetric alterations in OB in FEP patients. **(A)** Representative T1w (upper panel) and T2w (lower panel) images of the OB region of the same subject in the axial (left), coronal (middle), and sagittal (right) views. **(B)** Boxplot of total (left), right (middle), and left (right) OB volume between FEP patients and HCs. \*\*\**p* < 0.001.

Accordingly, we investigated the volumes of the total, right, and left OB in FEP patients compared to HCs, while controlling for confounding factors (see details in **Methods**). We observed a significant reduction in the volumes of the total (*p* = 2.12E-04), right (*p* = 5.14E-04), and left (*p* = 8.42E-04) OB in FEP patients compared with HCs (**Figure 3B**).

### Human study: pathological impact of local OE inflammation on the OB volume in FEP patients

Under the hypothesis that OE inflammation may have a pathological impact on the OB volume in FEP patients, we investigated an association between immune/inflammation-related DEGs (n = 9) (defined above, **Table S1**) and the OB volume in these patients. Partial correlation analysis controlling for confounding factors found a significant correlation between an increase in the total expression of immune/inflammation-related DEGs (measured by the first principal component of all 9 immune/inflammation-related DEGs) and a reduction in the right OB volume in FEP patients (total *p* = 0.05, right *p* = 0.03, and left *p* = 0.16) (**Figure 4A**). In contrast, such a correlation was not observed between the expression of 18 non-immune/inflammation-related DEGs and the OB volume (total *p* = 0.31, right *p* = 0.22, and left *p* = 0.49). We then examined HCs and found no significant correlation between 9 immune/inflammation-related DEGs and the OB volume (total *p* = 0.12, right *p* = 0.14, and left *p* = 0.13) in HCs (**Figure 4B**). Also, no correlation was observed between 18 non-immune/inflammation-related DEGs and the OB volume (total *p* = 0.59, right *p* = 0.55, and left *p* = 0.68) in HCs. These results suggest that immune/inflammation-related changes in ONCs/OE may have a more specific impact on the reduction of the OB volume in FEP patients.

**Figure 4.**
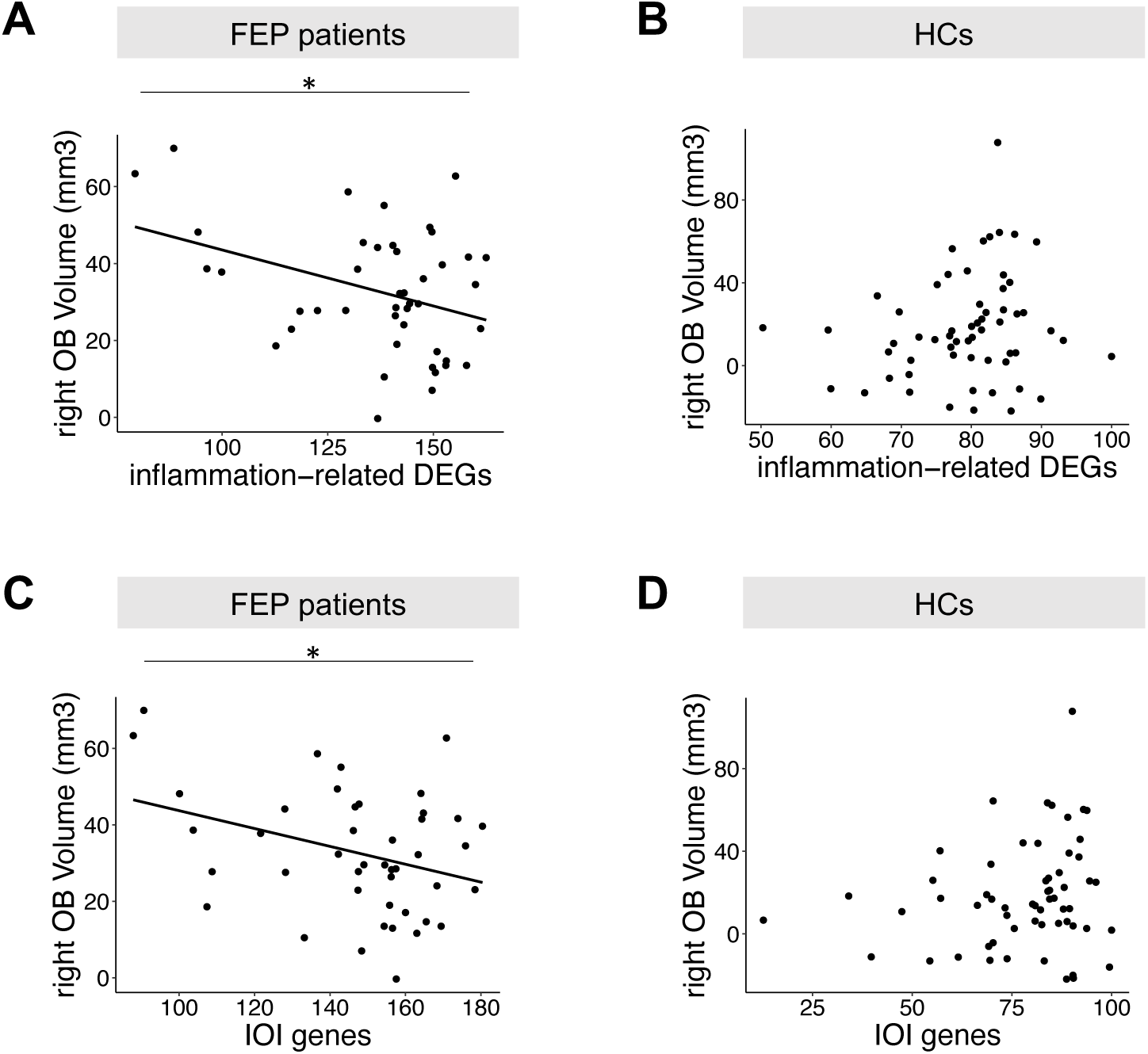
Correlation between OB volume and expression profiles in ONCs. The first principal component from PCA analysis was used to measure the total expression of a set of genes. A larger value means a higher expression level. **(A)** Dotplot of the expression of inflammation-related differentially expressed genes (DEGs) in ONCs *versus* right OB volume in FEP patients. **(B)** Dotplot of the expression of inflammation-related differentially expressed genes (DEGs) in ONCs *versus* right OB volume in human HCs. **(C)** Dotplot of the expression of human orthologs of the inflammation-related genes (IOI genes) in ONCs *versus* right OB volume in FEP patients. **(D)** Dotplot of the expression of human orthologs of the inflammation-related genes (IOI genes) in ONCs *versus* right OB volume in human HCs. \**p* < 0.05.

### Integration of mouse and human studies: pathological impact of local OE inflammation on the OB volume

To further examine the pathological impact of local OE inflammation on the OB volume, we conducted an integrative analysis using both mouse and human data. As described above, our past publication identified the IOI genes, a set of inflammation-related genes upregulated in the IOI mice compared with control mice^41^. Here, we examined the total expression of the human orthologs of the IOI genes in ONCs (measured by the first principal component of all 83 IOI genes) and their association with the OB volume in FEP patients. We observed a significant correlation between the increased expression of the IOI genes (n = 83) and a reduction of the right OB volume (total *p* = 0.09, right *p* = 0.04, and left *p* = 0.26) (**Figure 4C**). On the other hand, no significant correlations were observed between non-IOI genes (n = 14,534) and the OB volume in FEP patients (total *p* = 0.21, right *p* = 0.08, and left *p* = 0.54). We also examined HCs and found no significant correlations between the IOI genes and the OB volume (total *p* = 0.21, right *p* = 0.11, and left *p* = 0.44) in HCs (**Figure 4D**). No correlation was observed between non-IOI genes and the OB volume (total *p* = 0.60, right *p* = 0.56, and left *p* = 0.68) in HCs either. These results imply that the inflammatory molecular changes in ONCs are linked to a smaller OB volume, endorsing the notion that OE inflammation may impact the OB volume in FEP patients.

## Discussion

Smell deficits have been reproducibly reported in psychotic disorders, such as schizophrenia and related disorders^1–8^. Several studies have reported a reduced OB volume in these disorders^16,46,57–60^, but the sample size of these studies was relatively small. Multiple groups have prepared/enriched primary neuronal cells and stem cells from the OE via nasal biopsy from living subjects^30–32,35–39,57,61,62^. Molecular and cellular analyses of these cells from schizophrenia and related disorders have displayed deficits, indicating pathological alternations in the OE in these disorders^30–32,35–39,57,61,62^. Although changes associated with the olfactory system in patients with schizophrenia and related disorders are prominent and reproducibly reported from many groups^1,3,7,14,16,18,20,30,35,39,46,57–60,63^, their mechanistic link is unclear. To the best of our knowledge, no study has addressed the changes in OE and OB in a systematic manner, looking for the pathological relationship between the OE and OB. It is difficult to address the causal relationship between two distinct phenotypes, except in longitudinal studies with intervention. To fill these gaps, the present study combined the observations from both mice and humans. In human studies, we examined FEP patients with both brain imaging and molecular (next-generation sequencing) methodologies. We believe that the multimodal nature of the study design is a uniqueness and strength of the present study.

In the present study, by using the IOI model, we demonstrated a causal impact of chronic, local OE inflammation on OB volume and layer structure changes, reduced synaptic inputs to OB neurons. Furthermore, the IOI model displayed behavioral deficits relevant to negative symptoms (avolition) in parallel to smell deficits. In addition, we observed a significant alteration in the immune/inflammation-associated molecular signatures in OE-derived ONCs from FEP patients, implying that OE inflammation underlies the disease pathology. Concurrently, we also observed a significant reduction of the OB volume in FEP patients. In analogy to the OE-OB relationship in the IOI model, we looked for a potential link between the OE and OB in FEP patients via OE inflammatory changes, and demonstrated multiple lines of evidence that support such a link. First, we found a significant correlation between an increase in the expression of immune/inflammation-related DEGs and a reduction in the right OB volume in FEP patients, whereas a correlation was not observed between non-immune/inflammation-related DEGs and the OB volume in FEP patients. In contrast, no significant correlations were observed in HCs for either immune/inflammation-related or non-immune/inflammation-related DEGs. Second, we found a significant correlation between an increase in the expression of the IOI genes and a reduction in the right OB volume in FEP patients, whereas a correlation was not observed between non-IOI genes and the OB volume in FEP patients. Furthermore, no significant correlations were observed in HCs for either IOI or non-IOI genes. Taken together, we now report pathological molecular changes in the OE, reduced volume of the OB, and a possible link between OE inflammatory molecular changes and the reduced OB volume in FEP patients.

The associations between two types of inflammation-associated genes (immune/inflammation-related DEGs and IOI genes) and OB volume were observed only in FEP patients, but not in HCs. As described above, the OE is the most peripheral olfactory system located outside the cranium. As a result, the OE is directly exposed to several environmental risk factors, such as air pollution and viral infections in the upper respiratory tract. The difference in the immune/inflammation-related molecular signatures between FEP patients and HCs is likely a converged outcome of environmental factors and genetic risk factors associated with the disease. Furthermore, the OE-OB correlations only found in FEP patients, but not in HCs, may also imply the involvement of intrinsic (genetic) factors. This discussion is consistent with a clinical observation that, although a pool of patients with chronic rhinosinusitis (CRS) show cognitive changes^64–66^, all CRS patients do not show such mental manifestations. Together, at both OE pathology itself and the OE-OB relationship, disease-associated intrinsic factors may also play a role. In the future, with a much larger sample size, one may further test the impact of genetic factors on the outcome measures observed in the present study by utilizing polygenic risk scores. Although this is beyond the scope of the present study, the impact of the OE-OB pathological relationship on clinical manifestations in patients by considering the contributions of each environmental and genetic factor will be investigated as a significant research topic in the future.

In the present study, we also demonstrated the advantage of OE biopsy from living patients. Although we acknowledge the significance of postmortem tissues in research^67^, the cells/tissues obtained via biopsy reflect the condition of the subjects at the time of sampling. As shown in the present study, a multimodal assessment from living patients becomes available with OE biopsy. One potential drawback of OE biopsy is that it is technically difficult to maintain intact OE (neuroepithelium) at the quality for histopathological assessment in all biopsy cases. In contrast, we can enrich neuronal cells (e.g., ONCs) from any biopsied tissue, which facilitates scalability. Therefore, in the present study, we utilized ONCs to study pathological molecular signatures associated with OE inflammation in conjunction with the OB volume measurement in FEP patients. Nevertheless, to further validate and reinforce our observations, neuropathological and histological assessments of the intact neuroepithelium via biopsy may also be expected.

In summary, here we report OE pathological changes in immune/inflammatory pathways, OB structural changes, and their potential relationships in FEP patients. We believe that the compatibility of these patient data with those from the IOI mouse model reinforces the mechanistic link. Importantly, the electrophysiological data from OB neurons suggest that OE inflammatory changes can affect OB neurons that project to higher brain regions. This implies that a similar OE pathology found in FEP patients may also impact higher brain regions, such as the prefrontal cortex, via the olfactory-prefrontal circuits. The dysfunction of these circuits can underlie negative symptoms and some cognitive deficits that are known to associate with smell deficits. Taken together, the OE-OB data of the present study may be a basis for future studies that explore the implication of the olfactory system deficits in psychiatric manifestations in schizophrenia and related disorders.

Smell deficits are also reported in other neuropsychiatric conditions. Given that OE pathology is influenced by disease-specific genetic risk factors, the implication of OE pathology in smell deficits and neuropsychiatric manifestations may vary in each disease. Nevertheless, a cross-disease nature of smell deficits may also include common mechanistic changes shared among these diseases. It is known that SARS-CoV-2 infected patients frequently show smell deficits^68–70^. Post SARS-CoV-2 symptoms include psychosis, but several other neuropsychiatric conditions also occur^71,72^. Post SARS-CoV-2 neuropsychiatric manifestations may occur due to the viral-induced microvascular changes in the brain, but may also happen because of the impact from the OE-OB pathology. We should be careful and avoid a superficial link between the OE-OB pathology in schizophrenia and post SARS-CoV-2 neuropsychiatric conditions. Nevertheless, it is also possible that the mechanistic study of OE-OB pathology in schizophrenia and related disorders, as well as in other neuropsychiatric disorders, may provide valuable insight to post SARS-CoV-2 neuropsychiatric conditions.

## Data availability

The RNA-seq data and analysis scripts have been made publicly accessible on GitHub: https://github.com/KunYang99/Inflammation_ONC.

## Author contributions

The current research was designed by AS (Sawa). The analytic pipeline was designed by KY. The preclinical study regarding anatomical assessment and mouse behavioral test was designed by AK and carried out by YH. The preclinical study regarding patch clamp recording was designed by MM and carried out by JPB with the assistance of YW and YFZ. Both clinical and preclinical data were analyzed by KY, with the assistance of YH, JPB, MD, and SE. The anatomical assessment of the human olfactory bulb was supervised by JH and carried out by NP, LD, and AP. Nasal biopsy was designed and conducted by APL, with the assistance of AS (Smith). The olfactory neuronal cells were enriched from biopsied olfactory epithelium by KI. The clinical data interpretation was guided by AS (Sawa), JH, and VK. The manuscript was drafted by KY, YH, JPB, KI, MM, AK, and AS (Sawa). All authors contributed to the discussion of the results and have approved the final manuscript to be published.

## Acknowledgements

This study is supported by the National Institutes of Mental Health Grants MH-092443 (to AS), MH-094268 (to AS, AK), MH-105660 (to AS), MH-107730 (to AS), and MH-128765 (to AK); foundation grants from Stanley (to AS), RUSK/S-R (to AS), a NARSAD young investigator award from the Brain and Behavior Research Foundation (to AS, KY), NS-108452 (to JH, VK), AG-064093 (to JH, VK), DA-041208 (to AK), AG-065168 (to AK), DC-016106 (to AL), AI-132590 (to AL), DC-006213 (to MM), Kanae (YH), and Mitsui Sumitomo Insurance Welfare Foundation (YH). Figure. 1A and 2B were created with a graphical software provided by Biorender.com.

The authors wish to extend their gratitude to the participants in the current study. The authors thank Dr. Yu Miyahara and Ms. Vesna Tran for assisting mouse study, thank Ms. Yukiko Lema for research management and manuscript organization, and thank Dr. Melissa A Landek-Salgado for scientific and English editions.

## References

1 Cohen AS, Brown LA, Auster TL. Olfaction, ‘olfiction,’ and the schizophrenia-spectrum: an updated meta-analysis on identification and acuity. Schizophr Res 2012; 135: 152–157.

2 Ishizuka K, Tajinda K, Colantuoni C, Morita M, Winicki J, Le C et al. Negative symptoms of schizophrenia correlate with impairment on the University of Pennsylvania smell identification test. Neurosci Res 2010; 66: 106–110.

3 Kamath V, Lasutschinkow P, Ishizuka K, Sawa A. Olfactory Functioning in First-Episode Psychosis. Schizophr Bull 2018; 44: 672–680.

4 Kiparizoska S, Ikuta T. Disrupted Olfactory Integration in Schizophrenia: Functional Connectivity Study. Int J Neuropsychopharmacol 2017; 20: 740–746.

5 Kopala LC, Good K, Honer WG. Olfactory identification ability in pre- and postmenopausal women with schizophrenia. Biol Psychiatry 1995; 38: 57–63.

6 Malaspina D, Wray AD, Friedman JH, Amador X, Yale S, Hasan A et al. Odor discrimination deficits in schizophrenia: association with eye movement dysfunction. J Neuropsychiatry Clin Neurosci 1994; 6: 273–278.

7 Moberg PJ, Kamath V, Marchetto DM, Calkins ME, Doty RL, Hahn C-G et al. Meta-analysis of olfactory function in schizophrenia, first-degree family members, and youths at-risk for psychosis. Schizophr Bull 2014; 40: 50–59.

8 Turetsky BI, Hahn C-G, Borgmann-Winter K, Moberg PJ. Scents and nonsense: olfactory dysfunction in schizophrenia. Schizophr Bull 2009; 35: 1117–1131.

9 Chen X, Xu J, Li B, Guo W, Zhang J, Hu J. Olfactory impairment in first-episode schizophrenia: a case-control study, and sex dimorphism in the relationship between olfactory impairment and psychotic symptoms. BMC Psychiatry 2018; 18: 199.

10 Kamath V, Crawford J, DuBois S, Nucifora FC, Nestadt G, Sawa A et al. Contributions of olfactory and neuropsychological assessment to the diagnosis of first-episode schizophrenia. Neuropsychology 2019; 33: 203–211.

11 Kamath V, Turetsky BI, Calkins ME, Kohler CG, Conroy CG, Borgmann-Winter K et al. Olfactory processing in schizophrenia, non-ill first-degree family members, and young people at-risk for psychosis. World J Biol Psychiatry Off J World Fed Soc Biol Psychiatry 2014; 15: 209–218.

12 Good KP, Sullivan RL. Olfactory function in psychotic disorders: Insights from neuroimaging studies. World J Psychiatry 2015; 5: 210–221.

13 Cumming AG, Matthews NL, Park S. Olfactory identification and preference in bipolar disorder and schizophrenia. Eur Arch Psychiatry Clin Neurosci 2011; 261: 251–259.

14 Brewer WJ, Pantelis C, Anderson V, Velakoulis D, Singh B, Copolov DL et al. Stability of olfactory identification deficits in neuroleptic-naive patients with first-episode psychosis. Am J Psychiatry 2001; 158: 107–115.

15 Good KP, Tibbo P, Milliken H, Whitehorn D, Alexiadis M, Robertson N et al. An investigation of a possible relationship between olfactory identification deficits at first episode and four-year outcomes in patients with psychosis. Schizophr Res 2010; 124: 60–65.

16 Turetsky BI, Moberg PJ, Quarmley M, Dress E, Calkins ME, Ruparel K et al. Structural anomalies of the peripheral olfactory system in psychosis high-risk subjects. Schizophr Res 2018; 195: 197–205.

17 Corcoran C, Whitaker A, Coleman E, Fried J, Feldman J, Goudsmit N et al. Olfactory deficits, cognition and negative symptoms in early onset psychosis. Schizophr Res 2005; 80: 283–293.

18 Takahashi T, Nakamura M, Sasabayashi D, Komori Y, Higuchi Y, Nishikawa Y et al. Olfactory deficits in individuals at risk for psychosis and patients with schizophrenia: relationship with socio-cognitive functions and symptom severity. Eur Arch Psychiatry Clin Neurosci 2018; 268: 689–698.

19 Good KP, Whitehorn D, Rui Q, Milliken H, Kopala LC. Olfactory identification deficits in first-episode psychosis may predict patients at risk for persistent negative and disorganized or cognitive symptoms. Am J Psychiatry 2006; 163: 932–933.

20 Lin A, Brewer WJ, Yung AR, Nelson B, Pantelis C, Wood SJ. Olfactory identification deficits at identification as ultra-high risk for psychosis are associated with poor functional outcome. Schizophr Res 2015; 161: 156–162.

21 Mori K, Sakano H. How is the olfactory map formed and interpreted in the mammalian brain? Annu Rev Neurosci 2011; 34: 467–499.

22 Mori K, Sakano H. Olfactory Circuitry and Behavioral Decisions. Annu Rev Physiol 2021; 83: 231–256.

23 Hasegawa Y, Ma M, Sawa A, Lane AP, Kamiya A. Olfactory impairment in psychiatric disorders: Does nasal inflammation impact disease psychophysiology? Transl Psychiatry 2022; 12: 314.

24 Bhattarai JP, Etyemez S, Jaaro-Peled H, Janke E, Leon Tolosa UD, Kamiya A et al. Olfactory modulation of the medial prefrontal cortex circuitry: Implications for social cognition. Semin Cell Dev Biol 2022; 129: 31–39.

25 Kabbani N, Olds JL. Does COVID19 Infect the Brain? If So, Smokers Might Be at a Higher Risk. Mol Pharmacol 2020; 97: 351–353.

26 Zubair AS, McAlpine LS, Gardin T, Farhadian S, Kuruvilla DE, Spudich S. Neuropathogenesis and Neurologic Manifestations of the Coronaviruses in the Age of Coronavirus Disease 2019: A Review. JAMA Neurol 2020; 77: 1018–1027.

27 Gao Q, Xu Q, Guo X, Fan H, Zhu H. Particulate matter air pollution associated with hospital admissions for mental disorders: A time-series study in Beijing, China. Eur Psychiatry J Assoc Eur Psychiatr 2017; 44: 68–75.

28 Newbury JB, Arseneault L, Beevers S, Kitwiroon N, Roberts S, Pariante CM et al. Association of Air Pollution Exposure With Psychotic Experiences During Adolescence. JAMA Psychiatry 2019; 76: 614–623.

29 Pedersen CB, Raaschou-Nielsen O, Hertel O, Mortensen PB. Air pollution from traffic and schizophrenia risk. Schizophr Res 2004; 66: 83–85.

30 Evgrafov OV, Armoskus C, Wrobel BB, Spitsyna VN, Souaiaia T, Herstein JS et al. Gene Expression in Patient-Derived Neural Progenitors Implicates WNT5A Signaling in the Etiology of Schizophrenia. Biol Psychiatry 2020; 88: 236–247.

31 Rhie SK, Schreiner S, Witt H, Armoskus C, Lay FD, Camarena A et al. Using 3D epigenomic maps of primary olfactory neuronal cells from living individuals to understand gene regulation. Sci Adv 2018; 4: eaav8550.

32 Fan Y, Abrahamsen G, McGrath JJ, Mackay-Sim A. Altered cell cycle dynamics in schizophrenia. Biol Psychiatry 2012; 71: 129–135.

33 Féron F, Perry C, Hirning MH, McGrath J, Mackay-Sim A. Altered adhesion, proliferation and death in neural cultures from adults with schizophrenia. Schizophr Res 1999; 40: 211– 218.

34 English JA, Fan Y, Föcking M, Lopez LM, Hryniewiecka M, Wynne K et al. Reduced protein synthesis in schizophrenia patient-derived olfactory cells. Transl Psychiatry 2015; 5: e663.

35 Namkung H, Yukitake H, Fukudome D, Lee BJ, Tian M, Ursini G et al. The miR-124-AMPAR pathway connects polygenic risks with behavioral changes shared between schizophrenia and bipolar disorder. Neuron 2023; 111: 220–235.e9.

36 Jaaro-Peled H, Landek-Salgado MA, Cascella NG, Nucifora FC, Coughlin JM, Nestadt G et al. Sex-specific involvement of the Notch-JAG pathway in social recognition. Transl Psychiatry 2022; 12: 99.

37 Mihaljevic M, Lam M, Ayala-Grosso C, Davis-Batt F, Schretlen DJ, Ishizuka K et al. Olfactory neuronal cells as a promising tool to realize the ‘druggable genome’ approach for drug discovery in neuropsychiatric disorders. Front Neurosci 2022; 16: 1081124.

38 Takayanagi Y, Ishizuka K, Laursen TM, Yukitake H, Yang K, Cascella NG et al. From population to neuron: exploring common mediators for metabolic problems and mental illnesses. Mol Psychiatry 2021; 26: 3931–3942.

39 Kano S, Colantuoni C, Han F, Zhou Z, Yuan Q, Wilson A et al. Genome-wide profiling of multiple histone methylations in olfactory cells: further implications for cellular susceptibility to oxidative stress in schizophrenia. Mol Psychiatry 2013; 18: 740–742.

40 Hasegawa Y, Namkung H, Smith A, Sakamoto S, Zhu X, Ishizuka K et al. Causal impact of local inflammation in the nasal cavity on higher brain function and cognition. Neurosci Res 2021; 172: 110–115.

41 Chen M, Reed RR, Lane AP. Chronic Inflammation Directs an Olfactory Stem Cell Functional Switch from Neuroregeneration to Immune Defense. Cell Stem Cell 2019; 25: 501–513.e5.

42 Lane AP, Turner J, May L, Reed R. A genetic model of chronic rhinosinusitis-associated olfactory inflammation reveals reversible functional impairment and dramatic neuroepithelial reorganization. J Neurosci Off J Soc Neurosci 2010; 30: 2324–2329.

43 Saito A, Taniguchi Y, Rannals MD, Merfeld EB, Ballinger MD, Koga M et al. Early postnatal GABAA receptor modulation reverses deficits in neuronal maturation in a conditional neurodevelopmental mouse model of DISC1. Mol Psychiatry 2016; 21: 1449– 1459.

44 Dieterich A, Liu T, Samuels BA. Chronic non-discriminatory social defeat stress reduces effort-related motivated behaviors in male and female mice. Transl Psychiatry 2021; 11: 125.

45 Mihaljevic M, Lam M, Ayala-Grosso C, Davis-Batt F, Schretlen DJ, Ishizuka K et al. Olfactory neuronal cells as a promising tool to realize the “druggable genome” approach for drug discovery in neuropsychiatric disorders. Front Neurosci 2023; 16.https://www.frontiersin.org/articles/10.3389/fnins.2022.1081124.

46 Turetsky BI, Moberg PJ, Yousem DM, Doty RL, Arnold SE, Gur RE. Reduced olfactory bulb volume in patients with schizophrenia. Am J Psychiatry 2000; 157: 828–830.

47 Leucht S, Samara M, Heres S, Davis JM. Dose Equivalents for Antipsychotic Drugs: The DDD Method. Schizophr Bull 2016; 42 **Suppl 1**: S90–94.

48 Andrews S. FastQC: a quality control tool for high throughput sequence data. 2010.http://www.bioinformatics.babraham.ac.uk/projects/fastqc.

49 Martin M. Cutadapt removes adapter sequences from high-throughput sequencing reads. EMBnet.journal 2011; 17: 10–12.

50 Pertea M, Kim D, Pertea GM, Leek JT, Salzberg SL. Transcript-level expression analysis of RNA-seq experiments with HISAT, StringTie and Ballgown. Nat Protoc 2016; 11: 1650– 1667.

51 Love MI, Huber W, Anders S. Moderated estimation of fold change and dispersion for RNA-seq data with DESeq2. Genome Biol 2014; 15: 550.

52 Leek JT, Johnson WE, Parker HS, Jaffe AE, Storey JD. The sva package for removing batch effects and other unwanted variation in high-throughput experiments. Bioinformatics 2012; 28: 882–883.

53 Langfelder P, Horvath S. WGCNA: an R package for weighted correlation network analysis. BMC Bioinformatics 2008; 9: 559.

54 Reimand J, Arak T, Adler P, Kolberg L, Reisberg S, Peterson H et al. g:Profiler-a web server for functional interpretation of gene lists (2016 update). Nucleic Acids Res 2016; 44: W83–89.

55 Bhattarai JP, Schreck M, Moberly AH, Luo W, Ma M. Aversive Learning Increases Release Probability of Olfactory Sensory Neurons. Curr Biol CB 2020; 30: 31–41.e3.

56 Bressel OC, Khan M, Mombaerts P. Linear correlation between the number of olfactory sensory neurons expressing a given mouse odorant receptor gene and the total volume of the corresponding glomeruli in the olfactory bulb. J Comp Neurol 2016; 524: 199–209.

57 Yang K, Hua J, Etyemez S, Paez A, Prasad N, Ishizuka K et al. Volumetric alteration of olfactory bulb and immune-related molecular changes in olfactory epithelium in first episode psychosis patients. Schizophr Res 2021; 235: 9–11.

58 Asal N, Bayar Muluk N, Inal M, Şahan MH, Doğan A, Buturak SV. Olfactory bulbus volume and olfactory sulcus depth in psychotic patients and patients with anxiety disorder/depression. Eur Arch Oto-Rhino-Laryngol Off J Eur Fed Oto-Rhino-Laryngol Soc EUFOS Affil Ger Soc Oto-Rhino-Laryngol - Head Neck Surg 2018; 275: 3017–3024.

59 Nguyen AD, Pelavin PE, Shenton ME, Chilakamarri P, McCarley RW, Nestor PG et al. Olfactory sulcal depth and olfactory bulb volume in patients with schizophrenia: an MRI study. Brain Imaging Behav 2011; 5: 252–261.

60 Turetsky BI, Moberg PJ, Arnold SE, Doty RL, Gur RE. Low olfactory bulb volume in first-degree relatives of patients with schizophrenia. Am J Psychiatry 2003; 160: 703–708.

61 Rantanen LM, Bitar M, Lampinen R, Stewart R, Quek H, Oikari LE et al. An Alzheimer’s Disease Patient-Derived Olfactory Stem Cell Model Identifies Gene Expression Changes Associated with Cognition. Cells 2022; 11: 3258.

62 Lavoie J, Gassó Astorga P, Segal-Gavish H, Wu YC, Chung Y, Cascella NG et al. The Olfactory Neural Epithelium As a Tool in Neuroscience. Trends Mol Med 2017; 23: 100– 103.

63 Mor E, Kano S-I, Colantuoni C, Sawa A, Navon R, Shomron N. MicroRNA-382 expression is elevated in the olfactory neuroepithelium of schizophrenia patients. Neurobiol Dis 2013; 55: 1–10.

64 Jafari A, de Lima Xavier L, Bernstein JD, Simonyan K, Bleier BS. Association of Sinonasal Inflammation With Functional Brain Connectivity. JAMA Otolaryngol--Head Neck Surg 2021; 147: 534–543.

65 DeConde AS, Soler ZM. Chronic rhinosinusitis: Epidemiology and burden of disease. Am J Rhinol Allergy 2016; 30: 134–139.

66 Arslan F, Tasdemir S, Durmaz A, Tosun F. The effect of nasal polyposis related nasal obstruction on cognitive functions. Cogn Neurodyn 2018; 12: 385–390.

67 Arnold SE, Han LY, Moberg PJ, Turetsky BI, Gur RE, Trojanowski JQ et al. Dysregulation of olfactory receptor neuron lineage in schizophrenia. Arch Gen Psychiatry 2001; 58: 829– 835.

68 Tan BKJ, Han R, Zhao JJ, Tan NKW, Quah ESH, Tan CJ-W et al. Prognosis and persistence of smell and taste dysfunction in patients with covid-19: meta-analysis with parametric cure modelling of recovery curves. BMJ 2022; 378: e069503.

69 Liu N, Yang D, Zhang T, Sun J, Fu J, Li H. Systematic review and meta-analysis of olfactory and gustatory dysfunction in COVID-19. Int J Infect Dis IJID Off Publ Int Soc Infect Dis 2022; 117: 155–161.

70 Thaweethai T, Jolley SE, Karlson EW, Levitan EB, Levy B, McComsey GA et al. Development of a Definition of Postacute Sequelae of SARS-CoV-2 Infection. JAMA 2023; 329: 1934–1946.

71 Nakamura ZM, Nash RP, Laughon SL, Rosenstein DL. Neuropsychiatric Complications of COVID-19. Curr Psychiatry Rep 2021; 23: 25.

72 Premraj L, Kannapadi NV, Briggs J, Seal SM, Battaglini D, Fanning J, et al. Mid and long-term neurological and neuropsychiatric manifestations of post-COVID-19 syndrome: A meta-analysis. J Neurol Sci 2022; 434: 120162.

