## Supplemental Materials for "Inflammation-related pathology in the olfactory epithelium: its impact on the olfactory system in psychotic disorders"

**Table S1. Significant genes identified via differential expression analysis between healthy controls and first episode psychosis patients.**

Analysis results from DESeq2. Meaning of columns: baseMean, mean of normalized counts for all samples; log2FoldChange, log2 of fold change; lfcSE, standard error value; stat, Wald statistic; padj, adjusted p-value after multiple comparison correction; #Pubmed, the number of publications which mentioned the gene and the keyword "immune," "inflammation," or "inflammatory" in their titles or abstracts. Nine genes, categorized as immune/ inflammation-related based on the threshold of 20 publications, were highlighted in gray shadow.

| Gene | baseMean | log2FoldChange | lfcSE | stat | pvalue | padj | #Pubmed |
|---|---|---|---|---|---|---|---|
| TSLP | 536.91 | 0.67 | 0.16 | 4.08 | 4.59E-05 | 0.04 | 1673 |
| EGR2 | 630.97 | 1.18 | 0.30 | 3.99 | 6.74E-05 | 0.04 | 259 |
| AOC3 | 97.67 | -1.16 | 0.29 | -3.99 | 6.58E-05 | 0.04 | 174 |
| TRIB3 | 17921.37 | 0.59 | 0.15 | 4.02 | 5.79E-05 | 0.04 | 84 |
| CNTN1 | 298.30 | -1.37 | 0.34 | -4.01 | 6.08E-05 | 0.04 | 57 |
| RBFOX3 | 35.80 | 1.26 | 0.30 | 4.18 | 2.85E-05 | 0.03 | 49 |
| HSPA2 | 309.34 | -1.04 | 0.23 | -4.55 | 5.41E-06 | 0.02 | 44 |
| YWHAE | 45854.68 | -0.19 | 0.05 | -4.05 | 5.13E-05 | 0.04 | 38 |
| SIK1 | 3754.82 | 0.66 | 0.17 | 4.00 | 6.44E-05 | 0.04 | 28 |
| UBE2V1 | 39412.73 | -0.16 | 0.04 | -4.45 | 8.48E-06 | 0.02 | 18 |
| SOAT2 | 70.71 | -0.94 | 0.21 | -4.44 | 8.84E-06 | 0.02 | 11 |
| EIF4G2 | 132111.40 | -0.13 | 0.03 | -4.31 | 1.66E-05 | 0.03 | 11 |
| PAPPA2 | 1672.11 | 1.14 | 0.28 | 4.02 | 5.90E-05 | 0.04 | 11 |
| KLHDC7B | 945.63 | 1.03 | 0.26 | 3.91 | 9.11E-05 | 0.05 | 6 |
| DMGDH | 181.43 | 0.80 | 0.18 | 4.43 | 9.58E-06 | 0.02 | 5 |
| SGCG | 65.47 | 1.31 | 0.30 | 4.38 | 1.19E-05 | 0.02 | 5 |
| MLLT10 | 3972.80 | 0.17 | 0.04 | 4.27 | 1.92E-05 | 0.03 | 5 |
| ATP6V1C2 | 43.47 | 2.01 | 0.31 | 6.54 | 6.05E-11 | 8.43E-07 | 4 |
| HIST1H3D | 94.94 | -0.77 | 0.19 | -4.06 | 4.88E-05 | 0.04 | 4 |
| EFHC1 | 1758.49 | 0.27 | 0.07 | 4.02 | 5.85E-05 | 0.04 | 2 |
| TMEM189 | 39412.73 | -0.16 | 0.04 | -4.45 | 8.48E-06 | 0.02 | 1 |
| RTCB | 16622.00 | -0.10 | 0.02 | -3.99 | 6.68E-05 | 0.04 | 1 |
| TMEM189-UBE2V1 | 39412.73 | -0.16 | 0.04 | -4.45 | 8.48E-06 | 0.02 | 0 |
| GOLGA6L9 | 524.33 | 0.31 | 0.07 | 4.19 | 2.73E-05 | 0.03 | 0 |
| HIST1H2AD | 94.94 | -0.77 | 0.19 | -4.06 | 4.88E-05 | 0.04 | 0 |
| FAM161B | 1581.51 | 0.28 | 0.07 | 3.98 | 6.80E-05 | 0.04 | 0 |
| NPIPA5 | 11144.46 | 0.27 | 0.07 | 3.95 | 7.88E-05 | 0.04 | 0 |

**Table S2. Two-group comparison of expression levels of co-expression network modules between first-episode psychosis patients and healthy controls.**

Six modules were identified by the WGCNA method. Among 13,935 genes expressed in olfactory neuronal cells, 8488 were not assigned to any modules.

| module | size | pvalue | padj |
|---|---|---|---|
| I | 124 | 6.24E-05 | 3.74E-04 |
| II | 223 | 4.96E-03 | 0.01 |
| III | 89 | 0.34 | 0.65 |
| IV | 3332 | 0.43 | 0.65 |
| V | 1319 | 0.59 | 0.70 |
| VI | 360 | 0.80 | 0.80 |

### Table S3. PubMed search results for genes in module II.

| gene | #Pubmed |
|---|---|
| MMP2 | 5237 |
| VDR | 1807 |
| JAK1 | 1777 |
| ABCA1 | 1030 |
| MCC | 1019 |
| ADAM17 | 1008 |
| IL1R1 | 832 |
| IVD | 733 |
| CASP1 | 709 |
| STAT2 | 695 |
| LAMP1 | 658 |
| CD7 | 508 |
| C1S | 452 |
| NRP1 | 396 |
| GRN | 395 |
| CASP4 | 389 |
| C1R | 386 |
| LRP1 | 384 |
| IFI16 | 366 |
| LRRK2 | 348 |
| CTSK | 312 |
| STK11 | 297 |
| DCN | 271 |
| NR3C1 | 266 |
| ARNT | 252 |
| FGF7 | 216 |
| NMI | 177 |
| PRKCA | 163 |
| CTSS | 155 |
| NFKBIZ | 152 |
| ANGPTL2 | 146 |
| CTSA | 114 |
| JCHAIN | 106 |
| TMEM119 | 103 |
| SATB1 | 101 |
| SLIT2 | 100 |
| ATF1 | 89 |
| CLMP | 89 |
| SOCS5 | 89 |
| PBX1 | 85 |
| DTNB | 79 |
| FOXO4 | 68 |
| LY6E | 62 |
| NFATC4 | 59 |
| PDGFD | 57 |
| RYK | 50 |
| ELF1 | 46 |
| FOXC2 | 44 |
| TTC7A | 41 |
| RNASET2 | 37 |
| CUX1 | 34 |
| PLXNB2 | 33 |
| THRA | 33 |
| TRIM8 | 31 |
| SUFU | 30 |
| CTBP1 | 28 |
| RTP4 | 28 |
| UBE2L6 | 28 |
| DPYD | 27 |
| FBLN5 | 27 |
| PTPN13 | 26 |
| PLPP3 | 25 |
| SIK3 | 25 |
| TLE1 | 25 |
| BCAT1 | 24 |
| LPCAT2 | 23 |
| LTBP2 | 22 |
| STAG2 | 22 |
| TBX3 | 22 |
| FOXL1 | 21 |
| LRRFIP1 | 21 |
| CTPS1 | 20 |
| DCTD | 20 |
| PTPRG | 18 |
| SNX9 | 18 |
| SVEP1 | 18 |
| SECTM1 | 17 |
| TRPS1 | 17 |
| INPP4B | 16 |
| MXRA8 | 15 |
| AK3 | 14 |
| C1RL | 14 |
| LPAR6 | 14 |
| LY75 | 14 |
| GREM2 | 13 |
| OLFML3 | 13 |
| PTPRS | 13 |
| SIAE | 13 |
| BDH2 | 12 |
| CD99L2 | 12 |
| FKBP8 | 12 |
| RPS6KA2 | 11 |
| USP33 | 11 |
| ALDH3B1 | 10 |
| CD302 | 10 |
| CLCA2 | 10 |
| CMTR1 | 9 |
| PSD3 | 9 |
| SLC44A1 | 9 |
| TM9SF4 | 9 |
| VPS39 | 9 |
| PGAP3 | 8 |
| SLC9A9 | 8 |
| LIME1 | 7 |
| MICU1 | 7 |
| NEK9 | 7 |
| NME7 | 7 |
| RABEP1 | 7 |
| ZFR | 7 |
| ALPK2 | 6 |
| HECW2 | 6 |
| NIPBL | 6 |
| RGL1 | 6 |
| SEL1L3 | 6 |
| SH3BP4 | 6 |
| TBC1D5 | 6 |
| TOB2 | 6 |
| VPS11 | 6 |
| ZDHHC1 | 6 |
| BCAS3 | 5 |
| DDX60L | 5 |
| IFT140 | 5 |
| RAPGEF2 | 5 |
| SLC2A4RG | 5 |
| SMARCC2 | 5 |
| VPS9D1 | 5 |
| APBB1 | 4 |
| CARD17 | 4 |
| EXOC4 | 4 |
| LZTS2 | 4 |
| ZGPAT | 4 |
| ADGRL2 | 3 |
| ATP9B | 3 |
| C6orf89 | 3 |
| CALCOCO1 | 3 |
| CARD16 | 3 |
| FAXDC2 | 3 |
| HGSNAT | 3 |
| IPO9 | 3 |
| LRRN4CL | 3 |
| SLC9B2 | 3 |
| ST6GAL2 | 3 |
| STARD5 | 3 |
| TCTN1 | 3 |
| WDR11 | 3 |
| WDR6 | 3 |
| WRNIP1 | 3 |
| ZER1 | 3 |
| ANKRD23 | 2 |
| DCAF6 | 2 |
| EFHC1 | 2 |
| EVC2 | 2 |
| GLT8D2 | 2 |
| GRAMD4 | 2 |
| MIER1 | 2 |
| NXPE3 | 2 |
| P4HTM | 2 |
| PCDHGC3 | 2 |
| PLEKHA6 | 2 |
| RAD9B | 2 |
| RBM43 | 2 |
| TBC1D17 | 2 |
| TJAP1 | 2 |
| TMEM98 | 2 |
| TPK1 | 2 |
| WDR81 | 2 |
| ZADH2 | 2 |
| ADSSL1 | 1 |
| BTBD8 | 1 |
| CCDC140 | 1 |
| CFAP69 | 1 |
| DENND6B | 1 |
| EMG1 | 1 |
| GTF3C4 | 1 |
| KCNK15 | 1 |
| KCNS2 | 1 |
| PCDHGA1 | 1 |
| PCDHGA9 | 1 |
| PCDHGB1 | 1 |
| PCDHGB2 | 1 |
| PCDHGB7 | 1 |
| PCDHGC4 | 1 |
| PCDHGC5 | 1 |
| PDZRN3 | 1 |
| PPM1M | 1 |
| RBM44 | 1 |
| RRAGB | 1 |
| SHISA4 | 1 |
| SPG21 | 1 |
| TBC1D14 | 1 |
| WDR91 | 1 |
| ABHD15 | 0 |
| ANKRD39 | 0 |
| C3orf18 | 0 |
| CCDC148 | 0 |
| CDC26 | 0 |
| CTDSP2 | 0 |
| DISP2 | 0 |
| GCC1 | 0 |
| LY75-CD302 | 0 |
| MEIOC | 0 |
| NIPAL2 | 0 |
| PCDHGA10 | 0 |
| PCDHGA11 | 0 |
| PCDHGA12 | 0 |
| PCDHGA2 | 0 |
| PCDHGA3 | 0 |
| PCDHGA4 | 0 |
| PCDHGA5 | 0 |
| PCDHGA6 | 0 |
| PCDHGA7 | 0 |
| PCDHGA8 | 0 |
| PCDHGB3 | 0 |
| PCDHGB4 | 0 |
| PCDHGB5 | 0 |
| PCDHGB6 | 0 |
| PCMTD2 | 0 |
| RNF149 | 0 |
| SLC9B1 | 0 |
| SPATA6 | 0 |
| TMCO4 | 0 |
| TMEM140 | 0 |
| TSTD3 | 0 |

#### Figure legends

**Figure S1. No effect of chronic OE inflammation on body weight, locomotion, and anxiety.**

**(A)** Body weight. Control, n=9 (4 males, 5 females); IOI, n=7 (4 males, 3 females)

(**B)** Locomotion (left), percentage of time spent in central area (right) as indicated by the open field test. Control, n=9 (4 males, 5 females); IOI, n=7 (4 males, 3 females).

(**C**) The percentage of entries into the open arm as indicated by the elevated plus maze test. Control, n=19 (9 males, 10 females); IOI, n=17 (8 males, 9 females)

Data are presented as the mean ± s.e.m.

**Figure S1.**

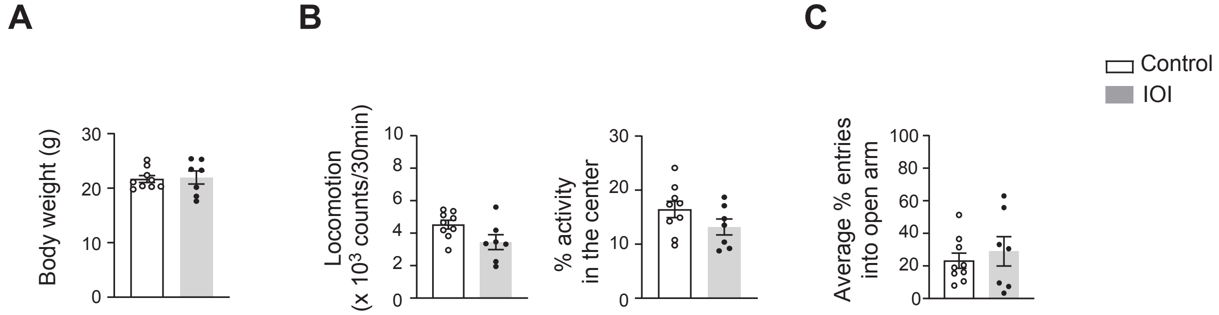
A
B
C
Control
IOI
Body weight (g)
0
10
20
30
Locomotion
(x 10^3 counts/30min)
0
2
4
6
8
10
% activity
in the center
0
10
20
30
Average % entries
into open arm
0
20
40
60
80
100
